# Increased Interstitial Flow and Elastic Lamina Degradation Precede Aortic Dissection

**DOI:** 10.64898/2026.07.09.737620

**Authors:** Shukei Sugita, Hiroomi Kaida, Yuki Hayashi, Aika Yamawaki-Ogata, Sakiko Nakamura, Yoshihiro Ujihara, Masanori Nakamura, Hideo Yokota, Yuji Narita

## Abstract

**Aims:** To elucidate the relationship between interstitial flow (IF) and structural changes in the aortic media during the development of aortic dissection (AD).

**Methods:** Apolipoprotein E-deficient (ApoE[–/–]) mice infused with angiotensin II (AngII) were used as an AD model, in which AD develops exclusively in the thoracoabdominal aorta but not in the thoracic aorta. To capture characteristics present prior to AD onset, the duration of AngII infusion was shortened to generate Pre-AD group. Normal C57BL/6 mice (Normal group) and ApoE(–/–) mice without AngII infusion (Control group) were also included for comparison. Thoracic and thoracoabdominal aortas were excised from all groups, and IF was measured *in vitro* in accordance with our previously established methods.

**Results:** IF velocity was generally smaller than 4 μm/s across all groups; however, velocities exceeding 4 μm/s was observed predominantly in the thoracoabdominal region of the Pre-AD group. Although mean IF velocity did not differ significantly among groups, the standard deviation differed significantly and was the highest in the thoracoabdominal region of the Pre-AD group. In this region, IF velocity tended to increase with more prolonged AngII administration, indicating acceleration of IF prior to AD onset. Three-dimensional internal structural microscopy revealed fragmentation of the elastic lamina (ELs) and a reduction in elastin density only in the thoracoabdominal region of the Pre-AD group.

**Conclusions:** Our findings suggest that increased IF and EL degradation occur in parallel and together contribute to the initiation of AD.

## 1 Introduction

Aortic dissection (AD) is a condition in which the aortic wall separates into two layers within the media, creating a double-lumen structure that extends longitudinally along the aorta. AD is a life-threatening disease, with a high mortality rate shortly after onset; approximately 50% of patients die within the first 24 hours^1^. Despite its severity, the underlying causes remain unclear^2^, and many aspects of its pathogenesis are still not fully understood. At present, it is generally thought that AD occurs when hemodynamic stress is imposed on a medial lesion, developing into a structurally weakened state over time^2^. Medial lesions include findings such as decreased elastic laminae (EL), reduced cross-linking fibers, and cystic medial degeneration^2^. Several studies^3–6^ on human AD have reported increased expression of matrix metalloproteinases (MMPs), especially MMP-2 and MMP-9, which degrade components of the vascular wall, suggesting the involvement of medial weakening. Hypertension constitutes a major hemodynamic risk factor for AD^7^, consistent with its high prevalence (71.8%) among patients with AD reported in a cardiovascular trial registry^8^. A large cohort study of about 30,000 people without aortic disease also showed that 86% of those who later developed AD were hypertensive at baseline^9^.

Hypertension increases the pressure gradient between the aortic lumen and the extravascular space and leads to an increase in interstitial flow (IF) velocity^10^. IF refers to the movement of fluid within the vessel wall from the luminal side toward the adventitial side. IF is thought to originate in the lumen, pass through the intima, then through fenestrations of approximately 1 µm in diameter within the internal elastic lamina^11^, and subsequently traverse the media and adventitia before exiting the vessel wall^12^. Numerical simulations have shown that elevated IF velocity increases shear stress on smooth muscle cells (SMCs) within the vessel wall^12–14^. Increased shear stress induces a phenotypic shift in SMCs from a contractile to a synthetic state^15, 16^, a transition that is known to occur in pathological conditions such as vascular injury^17^ and hypertension^18^. This synthetic phenotype is associated with enhanced proliferative and biosynthetic activity, including upregulated expression of MMP-2 and MMP-9^19^.

Taken together, these observations suggest that hypertension increases IF, leading to elevated shear stress on SMCs. This shear stress may induce phenotypic modulation of SMCs and upregulate MMP expression, thereby promoting loss of EL and degradation of elastin. Such changes could weaken the medial EL structure and potentially contribute to the onset of AD. However, if IF becomes faster specifically at sites where AD develops remains unknown. The present study aimed to elucidate the relationship between IF and structural changes in the medial layer during the development of AD. To this end, we examined IF and EL alterations before AD onset using an AD mouse model: Apolipoprotein E-deficient (ApoE[–/–]) mice infused with angiotensin II (AngII). The AngII infusion period was shortened to capture characteristics present prior to AD onset. In this model, AD develops exclusively in the thoracoabdominal aorta and not in the thoracic aorta^20, 21^. Accordingly, data from the AD model mice were compared between thoracic and thoracoabdominal segments. This approach enabled us to identify features specific to regions predisposed to AD development.

## 2 Methods

### 2.1 Sample preparation

All animal experiments were approved by the Institutional Review Board of Animal Care at Nagoya University (Approved; M220113-002, M230099-001) and Nagoya Institute of Technology (Approved; #2021006, #2022003) as well as the ARRIVE guidelines. Three types of mice were used in this study: Normal, Control, and Pre-AD groups. C57BL/6 mice (15.5–28.5 g; Japan SLC, Hamamatsu, Japan) were purchased for the Normal group. ApoE(–/–) C57BL/6 mice (19.0–26.5 g; Charles River Laboratories Japan, Yokohama, Japan) were used for the Control group. For the Pre-AD group, osmotic pumps (Alzet osmotic pump 2004, DURECT, Cupertino, CA, USA) were implanted into ApoE(–/–) mice (14.0–26.0 g), as described in previous studies^20–23^. AngII (05-23-0101, Calbiochem, Darmstadt, Germany) was administered at 750 ng/min/kg for 4–6 days. These mice develop AD primarily in the thoraco-abdominal region rather than in the thoracic region, as reported previously^20, 21^. In a preliminary experiment, we also tested an infusion rate of 1000 ng/min/kg for 4–5 days, following the method of Saraff et al^22^. However, the aortas became fragile and ruptured before inflation. Therefore, we reduced the AngII dose to induce a pre-AD condition without causing rupture. To check the heart rate and blood pressure, blood pressure and heart rate were measured using blood pressure measuring device (BP-98A-L, Softron, Tokyo, Japan) for a part of animals (*N* = 4–5).

The thoracic aorta was excised as described in previous studies^10, 24, 25^. Briefly, after the mice were euthanized in a CO_2_ chamber, thoracic aorta was exposed. Gentian violet dots were placed at 3-mm intervals on the surface of the aorta as *in vivo* length markers, and the intercostal arteries were cauterized. The aorta was then excised and kept in Krebs-Henseleit buffer (2.3 mM CaCl_2_, 115.3 mM NaCl, 4.6 mM KCl, 1.1 mM MgSO_4_, 22.1 mM NaHCO_3_, 1.1 mM KH_2_PO_4_, 7.8 mM glucose) until use.

### 2.2 Interstitial flow imaging

The interstitial flow was imaged as described in our previous study^10, 26^. Briefly, the excised tubular sample was pressurized using the experimental apparatus shown in Figure 1. Air pressure was regulated with an electropneumatic regulator connected to a pressure source (0.3 MPa), a digital-analogue/analogue-digital (DA/AD) converter, and a personal computer (PC). The regulated air pressure was applied to two reservoirs to convert air pressure into liquid pressure: one reservoir was filled with KH buffer, and the other was with 1.06 mM of uranine fluorescent dye solution (CI-45350, Tokyo Chemical Industry, Tokyo, Japan). The aortic sample was placed downstream of the reservoirs. Both ends of the sample were tied to hypodermic needles using suture threads, and the needles were fixed to a tissue bath. The sample was stretched longitudinally to reflect the *in vivo* state, using 3-mm interval markers. A pressure transducer positioned downstream of the sample measured the intraluminal pressure through a strain amplifier, the DA/AD converter, and the software (NI LabVIEW 2010) installed on the PC. The downstream side tubing was connected to an outlet bath via a three-way valve. The entire experimental system was maintained at 37°C using a warm circulation tank.

**Fig. 1.**
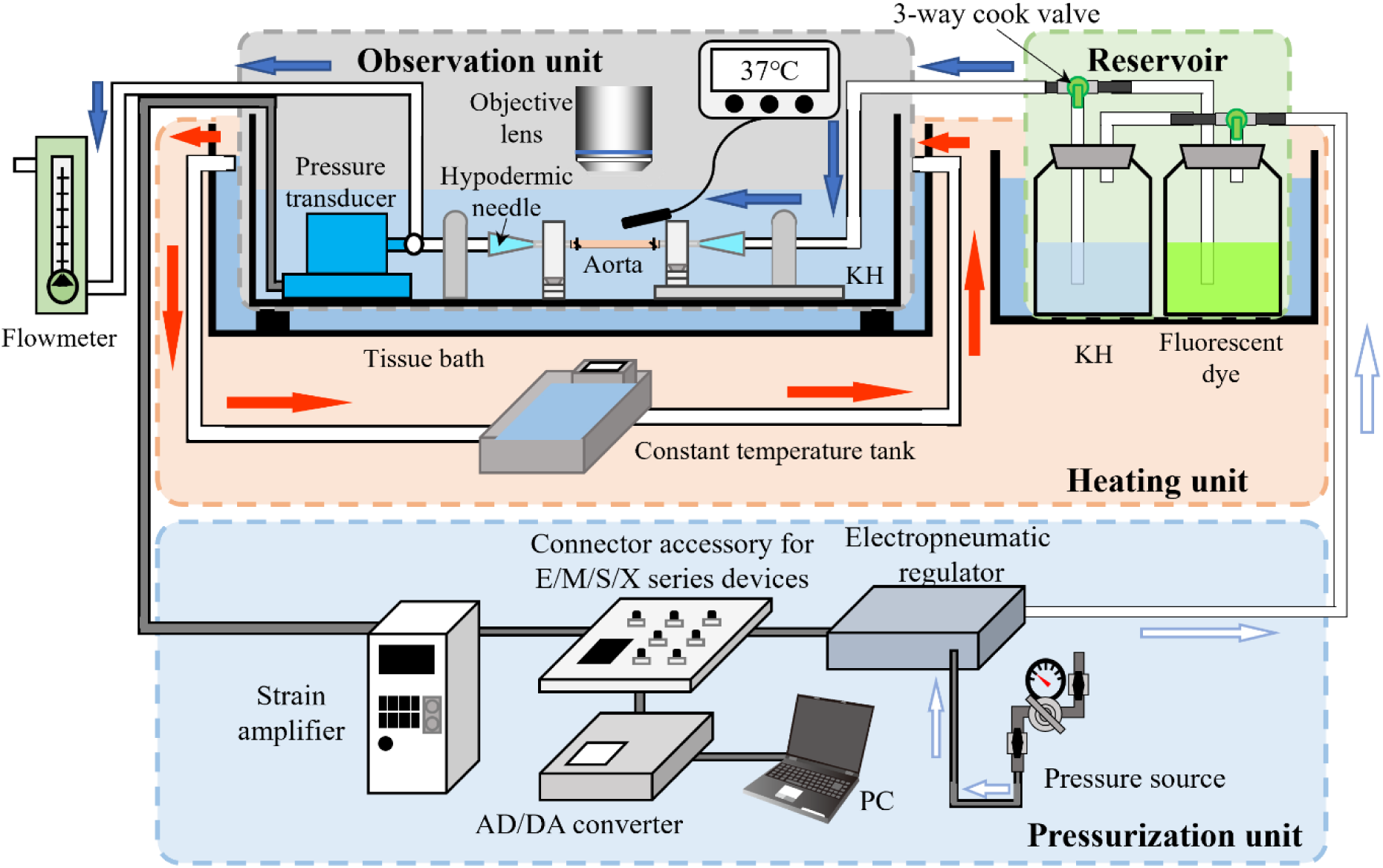
Schematic diagram of the interstitial flow measurement device. The apparatus consists of an observation unit (gray), a reservoir (green), an internal pressure loading unit (blue), and a heating unit (orange). The blue arrows indicate the direction of liquid flow, the white arrows indicate the direction of air flow, and the red arrows indicate the direction of hot water flow in the heating unit. KH, Krebs-Henseleit solution; DA/AD, digital-analogue/analogue-digital; PC, personal computer.

Fluorescent signals were captured using a two-photon microscope (FV1200MPE, Olympus, Tokyo, Japan). A Ti:sapphire laser (wavelength: 800 nm, output power: 2.0%) was directed onto the sample through a 60× objective lens (LUMPLFLN60XW, Olympus) and a V/G filter (FV-10-MRV/G, Olympus). To visualize the anatomical location of the aorta, autofluorescence from elastin was primarily detected using a V-channel bandpass filter (420–460 nm). Fluorescence from the dye solution was imaged using a G-channel bandpass filter (495–540 nm).

As the experimental protocol, the intraluminal surface of the specimen was pressurized, and fluorescence images were acquired. Image of 1×512 pixels (0.41×212 μm) were captured in the longitudinal *z* and circumferential *θ* plane, and 210–520 pixels (86.1–213.2 μm) were obtained in the radial *r* direction at 0.41-μm intervals. For some specimens, imaging was first performed around the celiac artery bifurcation. After each measurement, the fluorescent solution was washed out by perfusing KH buffer from the reservoir. This procedure was repeated 1–6 times at 80, 120, and 160 mmHg for each specimen by changing the observation site. All experiments were completed within 16 h after specimen harvest.

### 2.3 Deduction of the interstitial flow

Interstitial flow velocity was calculated as described previously^10, 26^. Briefly, all images were processed using ImageJ (v. 1.52a, National Institutes of Health, Bethesda, MD, USA). Image drift was corrected using the Image Stabilizer plugin. Because the elastic layers (ELs) were curved, images were converted from polar to Cartesian coordinates, and the time *t* series was linearly interpolated to a 1-s interval. The medial region between the internal EL (*r* = 0) and the external EL (*r* = *r*_EEL_) was extracted, circumferentially averaged, and resliced along the *r*-*t* plane to generate a kymograph. The kymograph was noise-reduced by compressing its size by *α* (1/10 in Sugita et al.^26^). Background subtraction and normalization set the left and right edges to 0 and 1, respectively. In the kymograph, the left and right boundary nodes were defined as the first points along both *r* = 0 and *r* = *r*_EEL_ where the normalized intensity reached *β* and 1-*β*, respectively (*β* = 0.10 from the Sugita et al.^26^). The region enclosed by these boundaries was used for analysis to exclude the non-equilibrated adventitial region and the saturated intimal region.

We assumed that the IF velocity *v* and diffusion coefficient *k* were constant along both the *r* and *t* axes. Thus, one-dimensional (1D) convection-diffusion equation was applied:

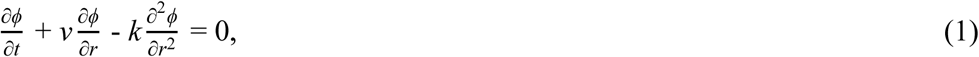

where *ϕ* represents the fluorescent dye concentration, corresponding to the fluorescence intensity in the experiment^10^. Discretized form of Eq. (1) is

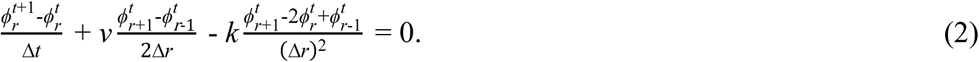

Here, *ϕ^t^* denotes the fluorescence intensity at coordinates (*t*, *r*) in the kymograph. The following terms can be computed directly from the kymograph:

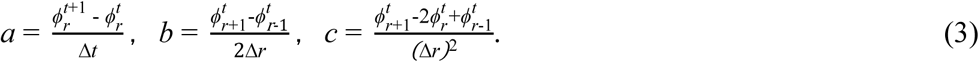

This yields a linear relationship of the form

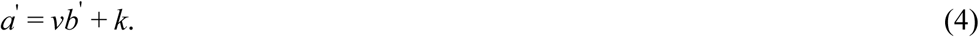

Here,

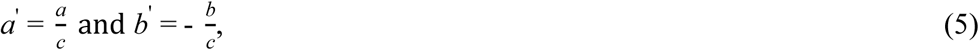

from which the IF velocity *v* was obtained by plotting *b’*-*a’* graph and fitting a linear regression. The slope *v* of the fitted linear regression represented the IF velocity. The significance of the correlation coefficient in the *b’*-*a’* plot was tested, and data showing no significant correlation were excluded from the analysis. Six to ten animals were included in each group.

### 2.4 Three-dimensional internal structure microscopy

Aortas from the three groups were imaged using three-dimensional (3D) internal structure microscopy (RMSS-002, RIKEN)^27^. Samples were prepared as described in the *Sample preparation* subsection, except that the adventitia was not removed. To preserve the straightened ELs, each sample was stretched to its *in vivo* length, and an intraluminal pressure of 20 mmHg was applied while the sample was immersed in 4% paraformaldehyde (168-23255, Fujifilm Wako Pure Chemical, Osaka, Japan) for 12 h to maintain its cylindrical shape.

The fixed samples were then sequentially immersed in 10%, 20%, and 30% sucrose in phosphate-buffered saline (PBS[–]) for 1 h each. After cryoprotection, the samples were embedded in Tissu Mount (Chiba Medical, Soka, Japan), frozen in liquid nitrogen, and stored at –80°C until imaging.

The specimen was observed using a 3D internal structure microscope. The blade rotation speed was set to 30 rpm, and the slicing thickness was adjusted to 2 µm. Images were acquired focusing on the EL in the *r-θ* cross-section of the vessel wall. An Apo Z ×1.5 objective lens (435281-9150-000, Carl Zeiss Microscopy, Oberkochen, Germany) and a Sona 4.2B-11 camera (Oxford Instruments, Abingdon-on-Thames, UK) were used. Laser light at wavelengths of 405, 488, and 637 nm was applied, and fluorescence was detected through a confocal scanner unit (CSU-W1, Yokogawa Electric, Tokyo, Japan) equipped with bandpass filters of 417–477 nm (405 channel), 475–525 nm (488 channel), and 645–685 nm (637 channel).

### 2.5 Image analysis of elastic laminas from the three dimensional internal structural image

Ten images were extracted from the 3D structural images at approximately equal longitudinal intervals (24–54 µm). In each image, the length of EL defects was measured. Defects were defined as partially missing or abnormally interrupted ELs (Fig. S1A). When a part of the vessel wall was outside the field of view or distorted, the image was halved; defect length was measured in one half and doubled to represent the full circumference. The total number of ELs in each image was counted, and the defect length was normalized by dividing by the number of ELs to obtain the defect length per EL, allowing comparison between the thoracic and thoracoabdominal regions.

To evaluate whether the reduction or structural disruption of the ELs occurred predominantly on the intimal or adventitial side, we quantified the radial distribution of EL density. Ten images used for defect length of EL were also used. Five circumferentially equally spaced straight lines were drawn radially so that they intersected the ELs perpendicularly (Fig. S1B and S1C), and intensities along each line were measured (Fig. S1D). Only high intensity data corresponding to ELs were left, and a linear regression line was fitted to these values (Fig. S1E). The intensity change ratio with respect to the *r*-axis for each image was defined and the averaged slopes of the five regression lines were evaluated.

### 2.6 Statistical method

Statistical analyses were performed using R version 4.6.0 (R Foundation for Statistical Computing, Vienna, Austria). For three mouse groups × two anatomical sites, a two-way analysis of variance (ANOVA) with heteroscedasticity-robust standard errors (HC3) was performed, as nonparametric alternatives do not adequately accommodate interaction effects. Post hoc pairwise comparisons were conducted using Tukey-adjusted tests based on estimated marginal means. For single-factor comparisons, nonparametric tests were applied due to violations of homogeneity of variance, including the Mann–Whitney U test for two-group comparisons and the Kruskal–Wallis test followed by the Steel–Dwass test for multiple-group comparisons. Homogeneity of variance was assessed using Bartlett’s test across multiple groups. Pearson’s correlation coefficients *r* were calculated, and their significance was assessed by testing the null hypothesis of no correlation. Data are presented as raw values as well as mean ± standard deviation (SD). Statistical significance was defined as *p* < 0.05. The number of observations is reported as *N* (number of mice per group) and *n* (number of measurements).

## 3 Results

### 3.1 Elevated Blood Pressure in the Pre-AD Group

Figure 2 summarizes heart rate, systolic blood pressure, diastolic blood pressure, and mean blood pressure in the Normal, Control, and Pre-AD groups. All measured parameters tended to be higher in the Pre-AD group than in other two groups. Significant differences from the Pre-AD group were observed for heart rate in the Control group, diastolic blood pressure in the Normal and Control groups, and mean blood pressure in the Control group.

**Fig. 2.**
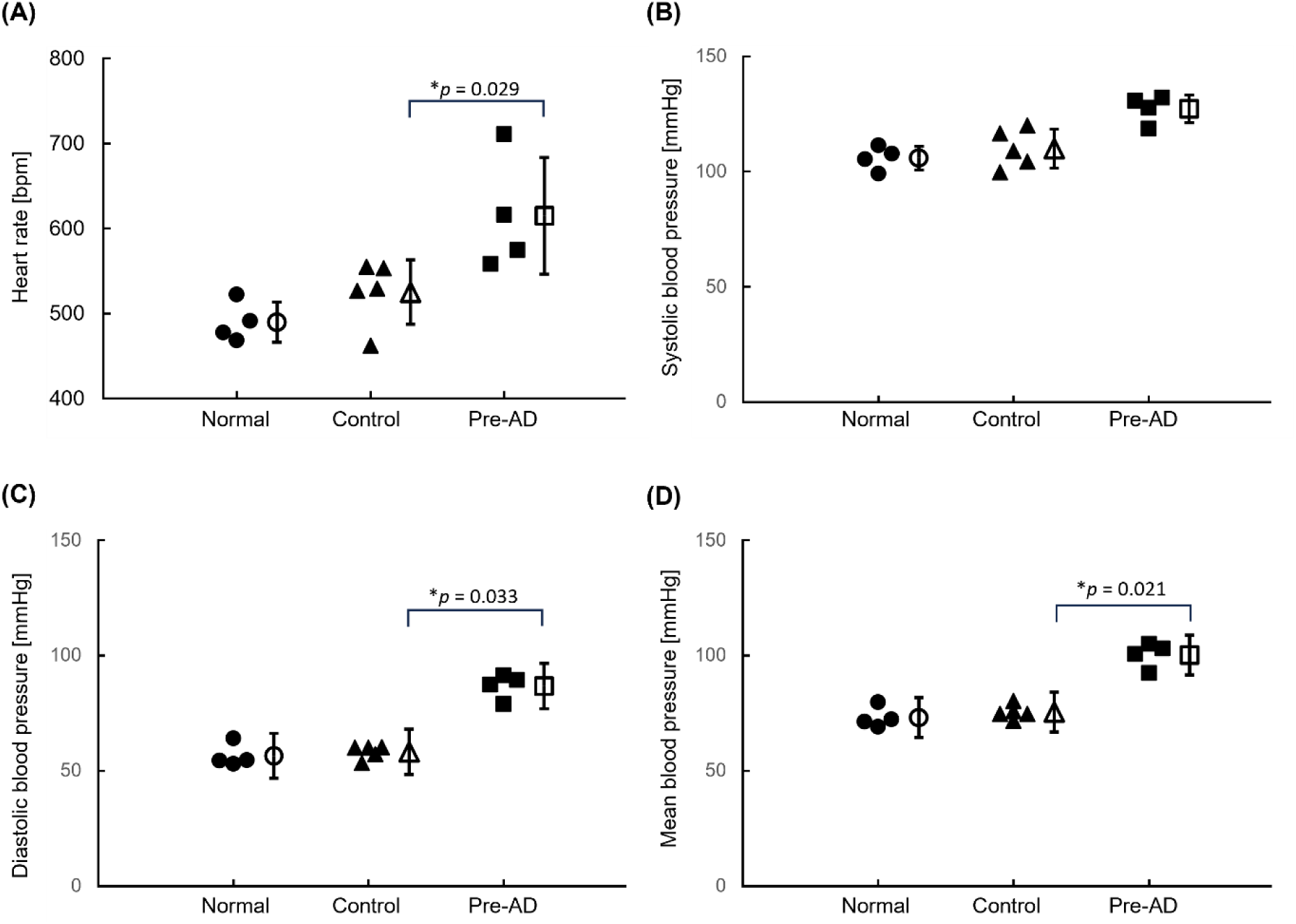
Physiological data of representative mice used in this study. (A) Heart rate, (B) systolic blood pressure, (C) diastolic blood pressure, and (D) mean blood pressure in the Control (*N* = 4), Normal (*N* = 5), and Pre-aortic dissection (Pre-AD, *N* = 4) groups. \**p* < 0.05 by Steel-Dwass test.

### 3.2 Structural Alterations of Elastic Lamellae in the Pre-AD Thoracoabdominal Aorta

Figure 3 shows representative fluorescent images of the *r*-*θ* plane from all groups during IF velocity measurements. In general, the ELs appeared curved, with relatively uniform spacing between adjacent ELs, except at the thoracoabdominal site in the Pre-AD group, where AD was expected to develop. In this region, the interlamellar distance was locally increased, suggesting radial separation of ELs. To evaluate the elongation of the distance between neighboring ELs, the maximum distance between ELs was measured in images obtained at 160 mmHg. Fig. 3G shows the distances between ELs for three mouse groups and two anatomical sites. Two-way ANOVA revealed significant main effects of group (*p* < 0.001) and site (*p* = 0.008), as well as a significant group × site interaction (*p* = 0.019). Post hoc analysis showed that the Pre-AD group exhibited significantly higher values than the Control and Normal groups in the thoracoabdominal region, whereas no significant differences were observed in the thoracic region. No significant differences between thoracic and thoracoabdominal regions were observed within each group. These results indicate that thoracoabdominal region in the Pre-AD group exhibits increased EL spacing. Furthermore, the aorta at the thoracoabdominal site in the Pre-AD group exhibited local loss of ELs (arrowhead in Fig. 3F). EL loss was observed in 54% (39/72) of images at this site in the Pre-AD group, while no EL loss was detected under any other condition.

**Fig. 3.**
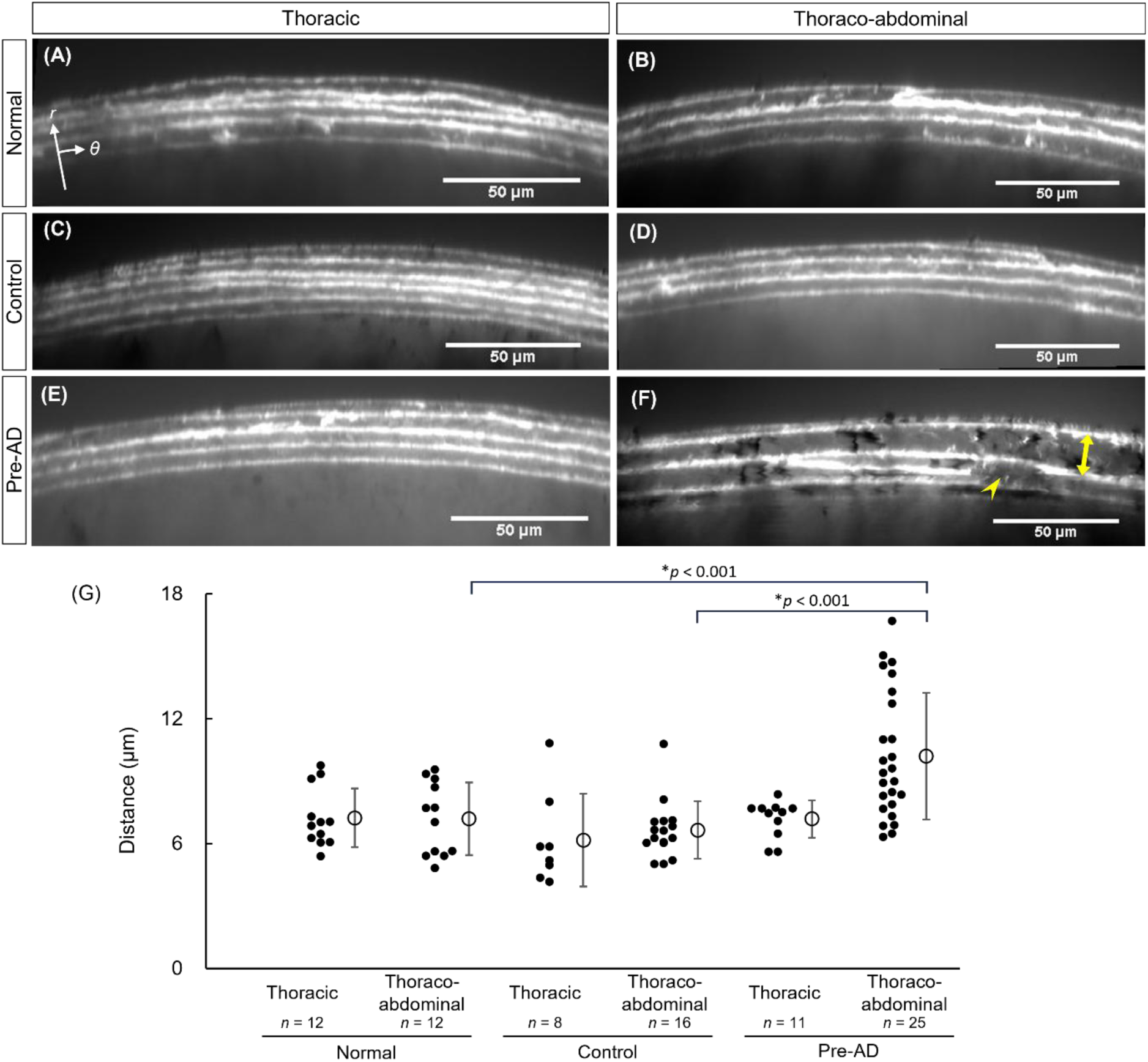
EL structure analyzed from images captured using a multiphoton microscope. (A–F) Representative images of elastin autofluorescence obtained from the (A, C, E) thoracic and (B, D, F) thoracoabdominal aorta of (A, B) Normal, (C, D) Control, and (E, F) Pre-AD mice. Yellow arrow indicates interlamellar elongation, and yellow arrowhead indicates a broken EL. (G) Maximum distance between neighboring ELs. AD, aortic dissection; EL, elastic lamina; *r*, radial; *θ*, circumferential axis. \**p* < 0.05 by Tukey-adjusted tests based on estimated marginal means.

### 3.3 Presence of Localized, Extremely High Interstitial Flow Velocity in the Thoracoabdominal Region in the Pre-AD Group

Movies 1–6 show the time-lapse changes in fluorescent dye concentration during IF velocity measurement in the six experimental conditions. From these movies, kymographs representing faster (Fig. S2A) and slower (Fig. S2B) IF cases were generated, and IF velocity was calculated using a 1D convection-diffusion equation. Figure 4 summarizes IF velocity in all groups. Across both anatomical sites and all three mouse groups, IF velocity was generally below 4 μm/s. Notably, IF velocities exceeding 4 μm/s were observed predominantly at the thoracoabdominal site in the Pre-AD group (Fig. 4F) across three different animals. Two-way ANOVA revealed a significant main effect of group at 80 mmHg, whereas no significant differences were detected at 120 or 160 mmHg. Post hoc Tukey’s HSD test indicated significant differences between the Normal and Pre-AD groups at 80 mmHg, with Pre-AD group exhibiting higher IF velocity. To further characterize differences among the six conditions, variability in IF velocity was assessed. Bartlett’s test demonstrated a significant difference in SD at 80 mmHg, but not at 120 or 160 mmHg. The SDs of IF velocity are summarized in Table 1. At 80 mmHg, the SD at the thoracoabdominal site in the Pre-AD group was 1.519 μm/s, exceeding those observed in all other conditions. Even at 160 mmHg, the SD at this site in the Pre-AD group tended to remain higher. Collectively, these findings suggest that localized regions of increased IF velocity are present in portions of the thoracoabdominal aorta in the Pre-AD group.

**Fig. 4.**
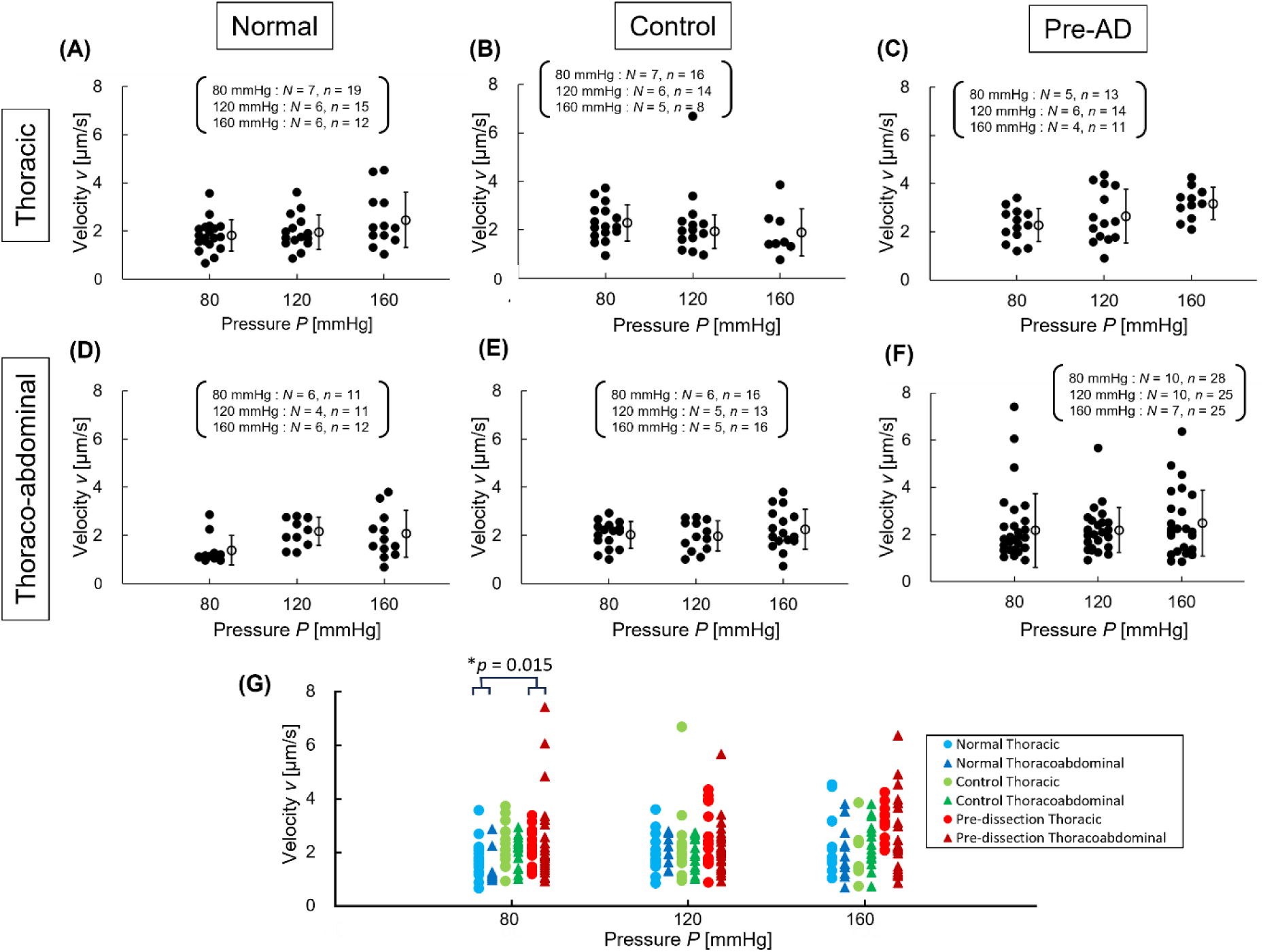
Interstitial flow velocity across the aortic walls. (A–F) Interstitial flow velocity of (A–C) thoracic and (D–F) thoraco-abdominal aorta in (A, D) Normal, (B, E) Control, and (C, F) Pre-AD groups. (G) Comparison between groups and aortic parts. AD, aortic dissection. \**p* < 0.05 by Tukey-adjusted tests based on estimated marginal means.

**Table 1.**
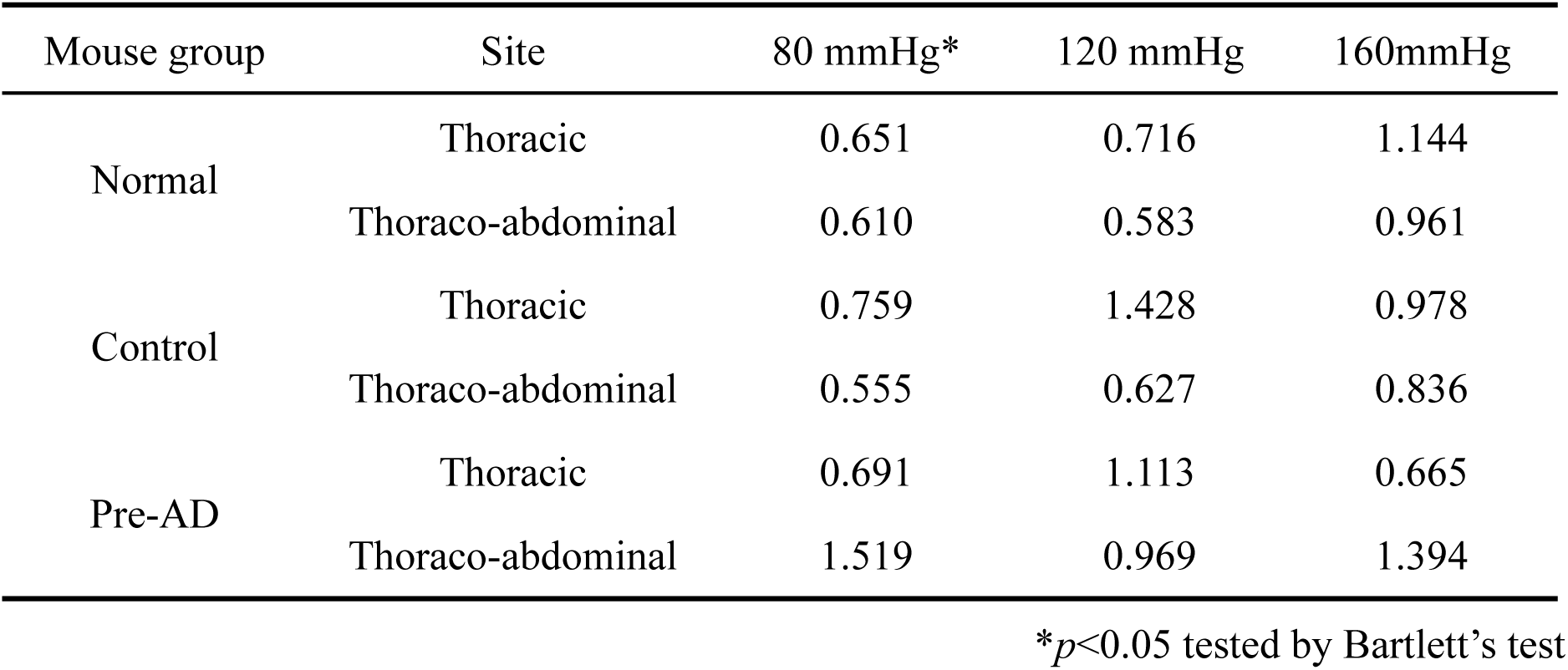
Standard deviation of IF velocity in each condition.

| Mouse group | Site | 80 mmHg* | 120 mmHg | 160mmHg |
| --- | --- | --- | --- | --- |
| Normal | Thoracic | 0.651 | 0.716 | 1.144 |
|  | Thoraco-abdominal | 0.610 | 0.583 | 0.961 |
| Control | Thoracic | 0.759 | 1.428 | 0.978 |
|  | Thoraco-abdominal | 0.555 | 0.627 | 0.836 |
| Pre-AD | Thoracic | 0.691 | 1.113 | 0.665 |
|  | Thoraco-abdominal | 1.519 | 0.969 | 1.394 |

\* $p < 0.05$ tested by Bartlett's test**Table 2** Average and Standard deviation (SD) of IF velocity in each duration of angiotensin II administration at the Pre-AD group in the thoraco-abdominal site.

To investigate potential factors underlying the locally elevated IF velocity, we examined the relationships between IF velocity and other parameters at the thoracoabdominal site in the Pre-AD group. The distance from the celiac artery bifurcation toward the downstream direction exhibited a weak positive trend with IF velocity (Fig. S3). However, regions of locally fast flow were also observed at variable positions, independent of this trend, indicating that longitudinal anatomical location does not strongly determine IF velocity. As shown in Fig. 3F, the thoracoabdominal site in the Pre-AD group exhibited localized EL loss and interlamellar elongation. Nevertheless, no significant differences in IF velocity were detected between images with and without local EL loss (Fig. S4A), or between images with and without interlamellar elongation (Fig. S4B). These results suggest that EL structural defects do not play a direct role in determining IF velocity.

In this study, the duration of AngII administration ranged from 4 to 6 days. Figure 5 plots the IF velocity against the duration of AngII administration. Although IF velocity tended to increase with longer AngII administration, no significant correlation was observed when analyzed at each pressure level. However, when all data points were analyzed collectively, IF velocity showed a significant correlation with the duration of AngII administration (*r* = 0.277, *p* = 0.014). Moreover, IF velocity was higher after 5 and 6 days of AngII administration than after 4 days. The Kruskal–Wallis test revealed significant differences at 120 and 160 mmHg of pressure. Post hoc Steel–Dwass testing showed significant differences between 4 and 5 days at 120 mmHg and between 4 and 6 days at 160 mmHg. These results indicate that a longer duration of AngII administration is associated with increased IF velocity. SD of IF velocity also tended to be higher at 5 and 6 days than 4 days of AngII administration period (Table 2). Bartlett’s test showed significant differences between administration days at 80, 120 mmHg, and all pressure data. These results indicate that SD of IF velocity was higher for longer AngII administration period. Thus, prolonged AngII exposure may modestly contribute to elevated IF velocity, suggesting that IF velocity may increase with the development of AD.

**Fig. 5.**
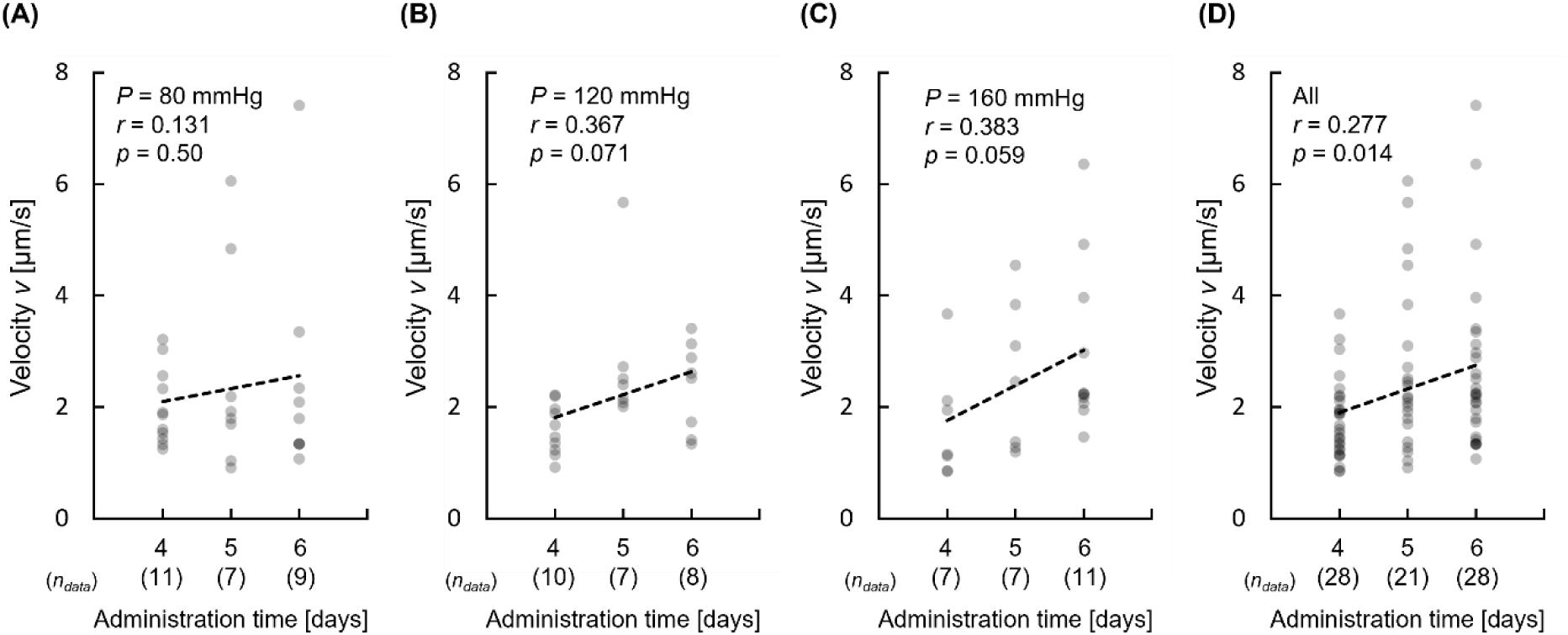
Effect of the duration of angiotensin II administration on interstitial flow velocity at intraluminal pressures of (A) 80, (B) 120, and (C) 160 mmHg. (D) Data from all pressure conditions combines. *n*_data_, Numbers of images analyzed. Pearson’s correlation coefficient (*r*) and corresponding *p* value are shown.

**Table 2.** Average and Standard deviation (SD) of IF velocity in each duration of angiotensin II administration at the Pre-AD group in the thoraco-abdominal site.

| Pressure | Average |  |  | SD |  |  |
| --- | --- | --- | --- | --- | --- | --- |
|  | 4 days | 5 days | 6 days | 4 days | 5 days | 6 days |
| 80 mmHg* | 2.01 | 2.56 | 2.46 | 0.69 | 1.86 | 1.99 |
| 120 mmHg* | 1.61 | 2.80 | 2.38 | 0.45 | 1.29 | 0.79 |
| 160 mmHg | 1.68 | 2.55 | 2.97 | 1.02 | 1.34 | 1.50 |
| All* | 1.78 | 2.63 | 2.64 | 0.71 | 1.47 | 1.50 |

\* $p < 0.05$ tested by Bartlett's test

### 3.4 Autofluorescence intensity of elastic lamina blurred at the thoracoabdominal site in Pre-AD group

Movies 7–13 and Figures 6A–6G show serial slices and a representative slice, respectively, of elastin autofluorescence in the *r*-*θ* plane obtained using 3D internal structure microscopy. ELs were clearly visualized and generally exhibited circular morphology, except at the thoracoabdominal site in the Pre-AD group. In this experiment, we incidentally observed an AD in one mouse that was intended to be prepared as a Pre-AD specimen.

**Fig. 6.**
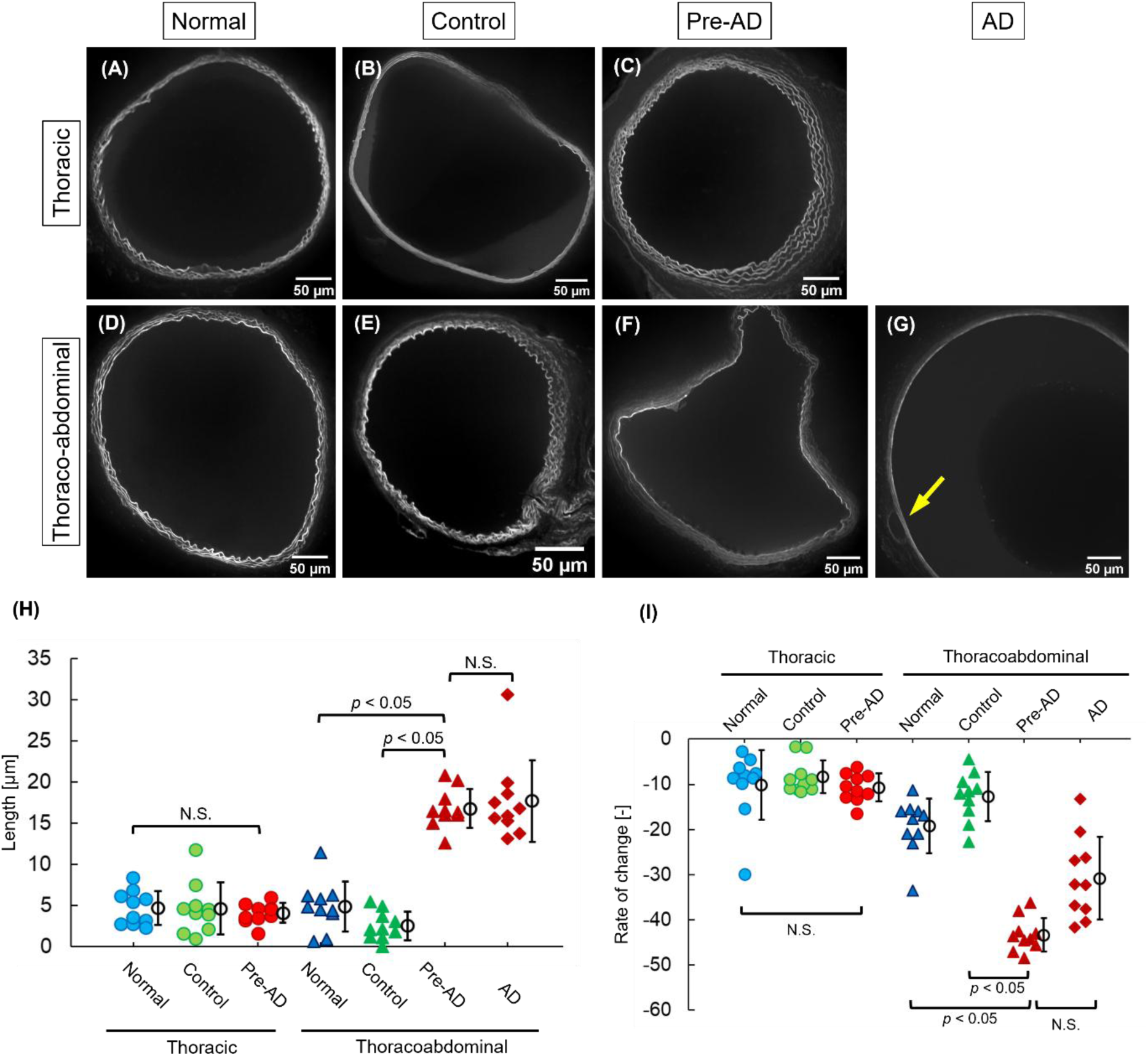
Changes in elastin analyzed using 3D internal structure microscope. The yellow arrow indicates an AD. (A–G) Representative cross-sectional images of the aorta at the (A–C) thoracic and (D–G) thoracoabdominal sites from (A, D) Normal, (B, E) Control, and (C, F, G) Pre-AD groups. Panel (G) shows the specimen with AD. See Movies 7–13 for sequential longitudinal images. (H) Length of broken ELs. (I) Rate changes in the autofluorescence intensity of ELs across the aortic wall from the intimal to adventitial side. AD, aortic dissection; EL, elastic laminas. \**p* < 0.05 by Tukey-adjusted tests based on estimated marginal means.

Figure 6H shows the length of fragmented ELs. Two-way repeated-measures ANOVA revealed significant main effects of group and anatomical position, as well as a significant interaction between these factors. Post hoc analyses indicated no significant differences at the thoracic site, whereas the thoracoabdominal site in the Pre-AD group exhibited significantly greater values. The AD specimen showed values comparable to those observed in the Pre-AD group.

Figure 6I illustrates the rate of change in EL density across the aortic wall along the *r*-axis. Two-way repeated-measures ANOVA demonstrated significant effects of group, anatomical position, and their interaction. Post hoc testing revealed no significant differences at the thoracic site; however, at the thoracoabdominal site, the Pre-AD group exhibited significantly lower values. The AD specimen again showed values similar to those of the Pre-AD group. Collectively, these results indicate that ELs at the thoracoabdominal site in the Pre-AD group were structurally degraded, particularly on the adventitial side.

## 4 Discussion

In this study, we identified distinct IF flow velocity and structural abnormalities in the thoracoabdominal aorta prior to the onset of AD. Notably, heterogeneity in IF velocity, reflected by an increased SD, was observed exclusively in the thoracoabdominal region of the Pre-AD group. Although group-wise mean IF velocity did not differ significantly, the presence of specimens exhibiting markedly elevated IF suggests that pathological alterations emerge locally and progressively at future AD sites rather than uniformly across the aorta. This interpretation is further supported by the observation that IF velocity tended to be higher with longer durations of AngII administration. Importantly, this localized increase in IF was accompanied by structural degradation of the EL, including EL fragmentation and reduced EL density, which were again restricted to the thoracoabdominal region in the Pre-AD group. These findings imply a mechanistic link between EL integrity and IF regulation. ELs play a critical role in maintaining radial structural continuity within the aortic wall; their degradation likely increases wall permeability and reduces resistance to transmural fluid transport. Consequently, disruption of EL architecture may facilitate abnormal IF penetration toward the adventitial side, as observed in the present study. Taken together, our findings support a hypothesis in which localized EL degradation precedes overt AD formation and contributes to abnormal IF dynamics within the aortic wall. Increased IF may, in turn, exacerbate structural weakening by promoting shear-related stress on the medial and adventitial layers, thereby creating a feed-forward loop that predisposes the thoracoabdominal aorta to dissection. These results highlight IF as a potential mechanistic mediator of early AD development.

In the thoracoabdominal region of the Pre-AD group, several specimens exhibited elevated IF velocity; however, no significant increase was observed at the group level. This finding indicates that only a subset of samples demonstrated abnormally rapid IF. One possible explanation for this heterogeneity is that ELs in these regions may undergo abrupt and localized structural failure. Such sudden disruption could eliminate local mechanical barriers within the aortic wall, thereby permitting interstitial fluid to traverse the tissue more rapidly. The ApoE(– /–) + AngII mouse model is known to exhibit substantial inter-individual variability in the timing of AD onset. While some mice develop AD within a few days following AngII infusion, others require more than ten days ^23, 28^. Consequently, specimens categorized within the Pre-AD group are likely to represent different stages along the continuum toward AD development. This variability in disease progression may manifest as heterogeneity in IF velocity, resulting in elevated IF in some specimens but not others, and ultimately contributing to the increased SD observed in the thoracoabdominal region of the Pre-AD group.

It remains unclear whether disruption of the EL leads to increased IF, or whether increased IF itself contributes to EL degradation. A straightforward interpretation is that EL fragmentation weakens the structural barrier of the aortic wall, thereby permitting IF velocity to increase. Supporting this view, studies using Fibulin-5 knockout mice and elastase-treated mice have reported increased hydraulic conductance—indicative of accelerated IF—suggesting that EL degradation precedes and promotes enhanced fluid transport^29^. Computational models of the aortic wall that explicitly incorporate IF dynamics further reinforce this concept. The simulations demonstrate that elevated blood pressure markedly increases radial tensile stress within the vessel wall^30^. Under conditions of EL degradation, the elevated blood pressure observed in the Pre-AD group may therefore amplify radial wall stress and facilitate AD initiation. Conversely, it is also plausible that increased IF precedes EL degradation and contributes causally to wall degeneration. The Pre-AD group generally exhibits higher blood pressure, which would be expected to drive faster IF^10^. Accelerated IF has been reported to promote a synthetic phenotype in SMCs^15^, a state associated with matrix remodeling. To the best of our knowledge, a direct relationship between elevated IF and MMP-2 or MMP-9 expression has not been reported; however, increased expression of MMP-1 has been demonstrated^31^, which may in turn accelerate EL degradation. Taken together, these findings suggest a bidirectional interplay between EL degradation and increased IF, potentially forming a self-reinforcing pathological loop during early AD development. At present, the causal hierarchy between these processes cannot be conclusively determined, and further studies will be required to elucidate their mechanistic relationship.

EL density decreased progressively toward the adventitial side. To our knowledge, no previous studies have explicitly reported a spatial decrease in EL density toward the adventitial side at sites predisposed to AD. However, several studies have presented histological images of the aorta from AngII-infused ApoE(–/–) mice in which the outer ELs appear less prominent, although this observation was not described or discussed in the accompanying text. Lu et al.^32^ presented Verhoeff-Van Gieson-stained sections in which ELs located farther from the lumen appeared more weakly stained. Similarly, Tham et al.^33^ provided Elastica Van Gieson-stained images showing that ELs in the outer layers of the aortic wall appeared lighter than those closer to the lumen. Although neither study explicitly described or quantified a decrease in EL density toward the adventitial side, these images are qualitatively consistent with the spatial trend observed in the present study. Nevertheless, these observations have been reported exclusively in AngII-infused ApoE(–/–) mouse models of AD. It therefore remains unclear whether a similar adventitial-side-dependent reduction in EL density occurs in human aortic tissue or in other models of AD.

The reason why EL density decreases more prominently toward the adventitial side remains unclear. One possible explanation is the influence of pro-inflammatory factors infiltrating from the adventitial side through the vasa vasorum. As shown by Daugherty et al.^20^, the pre-AD group develops a greater number of vasa vasorum compared with the normal group, suggesting that pro-inflammatory mediators may more readily penetrate into the medial layer. Once infiltrated, these inflammatory factors can alter intracellular signaling pathways in vascular SMCs^34, 35^, leading to phenotypic modulation from a contractile phenotype to an inflammatory phenotype^36^. Inflammatory SMCs are characterized by enhanced proliferative capacity and increased secretion of inflammatory mediators^37^. They have also been reported to upregulate several MMPs, including MMP-1 and MMP-3^38^, MMP-9^38, 39^, as well as MMP-2, MMP-13, and MMP-14^39^. Collectively, these findings suggest that infiltration of inflammatory mediators into the vessel wall may induce a phenotypic shift of SMCs from a contractile to an inflammatory state. Such a shift may lead to increased MMP activity, thereby contributing to elastin degradation.

## 5 Conclusion

To clarify the relationship between AD and IF, IF velocity was measured in aortic tissue prior to the onset of AD using an AngII-infused ApoE(–/–) mouse model. Markedly elevated IF velocity was observed predominantly in the thoracoabdominal region, where AD is predisposed to occur. IF velocity also tended to be higher with longer durations of AngII administration. Moreover, this region also exhibited structural weakening of ELs. Although the temporal sequence of these changes remains unclear, our findings strongly suggest that increased IF and EL degradation occur in parallel and together contribute to the initiation of AD.

## Non-standard Abbreviations and Acronyms

AD: aortic dissection
EL(s): elastic lamina(s)
MMP: matrix metalloproteinase
IF: interstitial flow
SMC(s): smooth muscle cell(s)
ApoE(–/–): apolipoprotein E-deficient
AngII: angiotensin II
DA/AD: digital-analogue/analogue-digital
PC: personal computer
KH: Krebs-Henseleit
1D: one-dimensional
3D: three-dimensional
SD: standard deviation

## Supplemental Materials

Figure S1–S4 Data Set Videos S1–S13

## Category of manuscript

Original Articles

## Novelty and Significance What Is Known?

- The mechanisms underlying the development of aortic dissection remain unclear.
- Hypertension is a well-established risk factor for aortic dissection.
- Increased intraluminal pressure within the aorta accelerates interstitial flow across the aortic wall.

## What New Information Does This Article Contribute?

- Extremely high interstitial flow velocities were observed predominantly in the thoracoabdominal region, where aortic dissection was expected to occur.
- Interstitial flow velocity remained elevated for longer durations of angiotensin II infusion, a condition required for the development of aortic dissection.
- Aortic structural alterations, including increased spacing between adjacent elastic laminae, elastin degradation, and fragmentation of elastic laminae, were observed at sites where extremely high interstitial flow velocities were recorded.
- Although degeneration of the aortic media is a hallmark of aortic dissection (AD), the biomechanical events that precede and potentially trigger this degeneration remain poorly understood. In particular, the role of interstitial flow (IF) in AD pathogenesis has not been investigated. Using an angiotensin II-induced mouse model, we demonstrate for the first time that localized acceleration of IF occurs before AD onset and is restricted to the thoracoabdominal aorta, a region susceptible to dissection. Importantly, elevated IF was spatially associated with fragmentation of elastic laminae and loss of elastin, suggesting a close interaction between altered transmural transport and medial degeneration during the earliest stage of disease development. These findings identify IF as a previously unrecognized biomechanical phenomenon associated with AD initiation. Rather than being a passive consequence of tissue damage, altered IF may participate in a positive-feedback process that promotes elastic lamina degradation and disease progression. This study provides a new mechanistic perspective on the initiation of AD and highlights IF as a potential biomechanical marker and therapeutic target for preventing dissection before catastrophic aortic failure occurs.

